# Structural Brain Alterations and Their Association with Cognitive Function and Symptoms in Attention-Deficit/Hyperactivity Disorder Families

**DOI:** 10.1101/863605

**Authors:** Wenhao Jiang, Kuaikuai Duan, Kelly Rootes-Murdy, Pieter J. Hoekstra, Catharina Hartman, Jaap Oosterlaan, Dirk Heslenfeld, Barbara Franke, Jan Buitelaar, Alejandro Arias-Vasquez, Jingyu Liu, Jessica A. Turner

**Affiliations:** Department of Psychology, Georgia State University, USA; School of Electrical and Computer Engineering, Georgia Institute of Technology, USA; University of Groningen, University Medical Center Groningen, Department of Psychiatry, Groningen, The Netherlands; Department of Clinical Neuropsychology, VU University Amsterdam, Amsterdam, The Netherlands; Department of Human Genetics, Donders Institute for Brain, Cognition and Behaviour, Radboud University Medical Center, Nijmegen, The Netherlands; Department of Psychiatry, Donders Institute for Brain, Cognition and Behaviour, Radboud University Medical Center, Nijmegen, The Netherlands; Department of Cognitive Neuroscience, Donders Institute for Brain, Cognition and Behaviour, Radboud University Medical Center, Nijmegen, The Netherlands; Department of Computer Science, TReNDS center, Georgia State University, Atlanta, USA; Neuroscience Institute, Georgia State University, USA

**Keywords:** ADHD, Independent component analysis, Cerebellum, Insula, Inattention

## Abstract

Gray matter disruptions have been found consistently in Attention-deficit/Hyperactivity Disorder (ADHD). The organization of these alterations into brain structural networks remains largely unexplored. We investigated 508 participants (281 males) with ADHD (N = 210), their unaffected siblings (N = 108), individuals with subthreshold ADHD (N = 49), and unrelated healthy controls (N = 141) with an age range from 7 – 18 years old from 336 families in the Dutch NeuroIMAGE project. Source based morphometry was used to examine structural brain network alterations and their association with symptoms and cognitive performance. Two networks showed significant reductions in individuals with ADHD compared to unrelated healthy controls after False Discovery Rate correction. Component A, mainly located in bilateral Crus I, showed a case/control difference with sub-clinical cases being intermediate between cases and controls. The unaffected siblings were similar to controls. After correcting for IQ and medication status, component A showed a negative correlation with inattention symptoms across the entire sample. Component B included a maximum cluster in the bilateral insula, where unaffected siblings, similar to cases, showed significantly reduced loadings compared to controls; but no relationship with individual symptoms or cognitive measures was found for component B. This multivariate approach suggests that areas reflecting genetic liability within ADHD are partly separate from those areas modulating symptom severity.

## 1. Introduction

Attention-deficit/Hyperactivity Disorder (ADHD) is a heritable neurodevelopmental disorder characterized by inattention and/or hyperactivity and impulsivity (Polanczyk and Rohde, 2007). Heritability of the disorder is estimated to be around 75%, with siblings of ADHD cases having a four times higher risk of developing ADHD than the general population (Biederman et al., 1990; Wolfers et al., 2016). Family and genetic factors influence both ADHD risk and brain structures (Demontis et al., 2017; Faraone et al., 2005; Klein et al., 2017; Peper et al., 2007). Unaffected siblings of ADHD cases show altered brain phenotypes, often at intermediate levels between cases and controls (Bralten et al., 2016), suggesting endophenotype qualities of such traits (Durston et al., 2006; Gottesman and Gould, 2003; Greven et al., 2015; Hart et al., 2014; van Rooij et al., 2015). However, the exact brain mechanisms behind such familial effects and their potential association with the symptoms and cognitive deficits relevant to ADHD are still unclear.

Structural brain alterations associated with ADHD have been reported consistently across cortical and subcortical regions (Ellison-Wright et al., 2008; Halperin and Schulz, 2006; Hoogman et al., 2017). Previous studies have demonstrated significant brain developmental delay and 3-5% smaller whole brain volume in individuals with ADHD compared to healthy controls (Castellanos et al., 1996; Castellanos et al., 2002; Greven et al., 2015). Meta-analysis studies have revealed ADHD-related brain abnormalities consistently in the caudate and basal ganglia, including the globus pallidus and putamen (Frodl and Skokauskas, 2012; Rogers and De Brito, 2016; Valera et al., 2007). Structural alterations of fronto-striatal-parietal pathways (Dickstein et al., 2006), cerebellum (Valera et al., 2007), anterior cingulate cortex (Narr et al., 2009; Norman et al., 2016), and several other brain regions have also been reported in relation to ADHD. Voxel-based morphometry (VBM) analyses of unaffected siblings of ADHD cases have identified alterations in the prefrontal cortex, medial and orbitofrontal cortex, fronto-occipital regions, and cingulate regions compared to healthy controls (Bralten et al., 2016; Durston et al., 2004; Pironti et al., 2014).

The brain regions listed above are involved in different disorders and cognitive deficits (Norman et al., 2016). It is well-known that brain regions do not act in isolation to support brain functions and different approaches have been used to define brain networks as possible “units” underlying function (Jadidian et al., 2015). A recent 2018 study employed source-based morphometry (SBM) (Xu et al., 2009), a data–driven decomposition approach, to extract brain network as potential markers in adult ADHD (Duan et al., 2018). Inspired by Duan and colleagues, we investigated multivariate structural brain network alterations in children with ADHD, compared to their unaffected siblings and unrelated controls. We subsequently examined the associations between the observed altered structural brain networks and major symptom domains and cognitive functions (namely, working memory and inhibition) while controlling for family structure. Thus, with family structure taken into account, we aimed to distinguish multivariate brain networks markers potentially associated with ADHD.

## 2. Material and methods

### 2.1. Participants

This study included 508 participants from 336 families from the NeuroIMAGE project (von Rhein et al., 2015). In this longitudinal study, families with an individual with ADHD were recruited, along with healthy control families. All participants provided written consent; detailed recruitment procedures, ethical approval, inclusion and exclusion criteria, as well as assessment information can be found in the previous description paper (von Rhein et al., 2015). Participants with ADHD were diagnosed according to the Diagnostic and Statistical Manual of Mental Disorders, 4th Edition (DSM-IV) (American Psychiatric Association, 1994). All participants had an IQ equal or greater than 70 as assessed by the Wechsler Adult Intelligence Scale (WAIS), and the absence of autism, epilepsy, learning difficulties, brain disorders, and genetic disorders mimicking symptoms of ADHD were confirmed (von Rhein et al., 2015).

Inattention and hyperactivity/impulsivity were assessed based on the Dutch translation of the Schedule for Affective Disorders Schizophrenia—present and lifetime version (K-SADS) and Conners’ Teacher Rating Scale −1997 Revised Version: Long Form, DSM-IV Symptoms Scale (CTRSR:L). The symptom counts from K-SADS and CTRSR:L were combined together, and the range of symptom scores was between 0 and 9 for each domain with a higher score indicating more severe symptoms. Participants were grouped into one of four categories: Those with ADHD, unaffected siblings, individuals with subthreshold ADHD, and non-ADHD controls (excluding siblings of ADHD cases), to follow groupings used in previous studies (von Rhein et al., 2015). Sub-clinical individuals had scores > 2 but < 5 in either domain while non-ADHD controls had ≤ 2 in either domain (von Rhein et al., 2015).

### 2.2. Neurocognitive assessments

Cognitive assessments available in the NeuroIMAGE sample included in the current study were evaluations of working memory and inhibition, two of the common cognitive deficits seen in ADHD (Alderson et al., 2013; Tarver et al., 2014). In the WAIS Digit Span test, maximum forward and backward scores were included (Wechsler, 2000). The average accuracy scores from Visuo-spatial Working Memory test were also included (Nutley et al., 2010). In a stop task, the stop signal reaction time, the deviation of reaction time from the mean, and total numbers of commission and omission error were evaluated to measure inhibition (Logan et al., 1984).

### 2.3. Image Acquisition and Processing

T1-weighted images were acquired from two sites with comparable 1.5T scanners (Sonata and Avanto; Siemens) with the following settings: a voxel size of 1×1×1mm^3^, TI 1000 ms, TR 2730 ms, TE 2.95 ms, field of view 256 mm, and 176 sagittal slices. Two independent raters applied a 4-point quality assurance scale to all the scans (1 = good; 2 = useable; 3 = poor; 4 = very poor), and only those images rated as “good” were included in the following analyses (von Rhein et al., 2015).

All images were segmented using Statistical Parametric Mapping 12 (SPM12, http://www.fil.ion.ucl.ac.uk/spm/software/spm12/), with age-specific templates generated by Template-O-Matic toolbox using the matched pairs approach (Wilke et al., 2008). Gray matter (GM) volumetric data were normalized to the Montreal Neurological Institute (MNI) template, modulated, and smoothed with a 6×6×6 mm^3^ Gaussian kernel. We performed a correlation analysis between the images and the original MNI template and all images showed greater than 0.8 correlation (mean *r* = 0.98). In addition, we applied a gray matter mask to the images which excluded voxels that had less the 20% gray matter. To remove possible confounding effects of gender and site, we performed a voxel-wise linear regression model with all images. Then we tested age and age^2 in following models to avoid driving the results of the voxel-wise linear regression.

### 2.4. Image decomposition and analysis

We utilized the SBM module of the GIFT Toolbox to perform independent component analysis (ICA) and component estimation using the minimum description length algorithm (Rissanen, 1978) From the infomax ICA algorithm with ICASSO within the GIFT toolbox, twenty distinct components were produced (Bell and Sejnowski, 1995; Xu et al., 2009). ICASSO (Himberg et al., 2004) with 10 ICA runs ensured the stability of components. The loading coefficients for each component and subject from these ICA results were the dependent variables in the following analyses, which reflect individual gray matter volume of each component. We included medication use as a binary variable (only in ADHD participants) and tested the quadratic effect of age (age^2) for all the 20 components. Further analyses included only the components showing significant medication or age^2 effects. Co-diagnoses of depression or anxiety were not included because only one participant was identified as having anxiety and eight participants were identified as having depression.

We applied a linear mixed model using family as a random effect with other variables as fixed effects (grouping, age, age^2, and medication when applicable), to detect components that differed between persons with ADHD and healthy controls. We applied the linear model using MATLAB function of fitlmematrix (MATLAB and Statistics Toolbox Release 2013a, The MathWorks, Inc., Natick, Massachusetts, United States.). We only included GM components showing significant case/control differences after a false discovery rate (FDR) correction of q < 0.05 in further association analyses for all participants. In addition, we calculated a moving average of loading coefficients from the significant components. The moving averages were based on the three-years-age bins, moving one year up every step. Those age bins with less than 6 participants were not included.

Finally, we conducted association analyses between component loadings and cognitive performance along with symptom scores with family random effects. Considering the potential confound of IQ, the final fixed effects in this model included IQ alongside with component loadings, age, gender, and medication.

## 3. Results

In decomposing the gray matter data of 508 children and adolescents including 210 ADHD patients, 108 unaffected siblings, 49 individuals with subthreshold ADHD, and 141 unrelated healthy controls (detailed demographic information is presented in Table 1), 20 brain components were generated and their ICASSO stability indices were all large (> 0.97). See supplemental Figure 1 for representations of all components.

**Figure 1.**
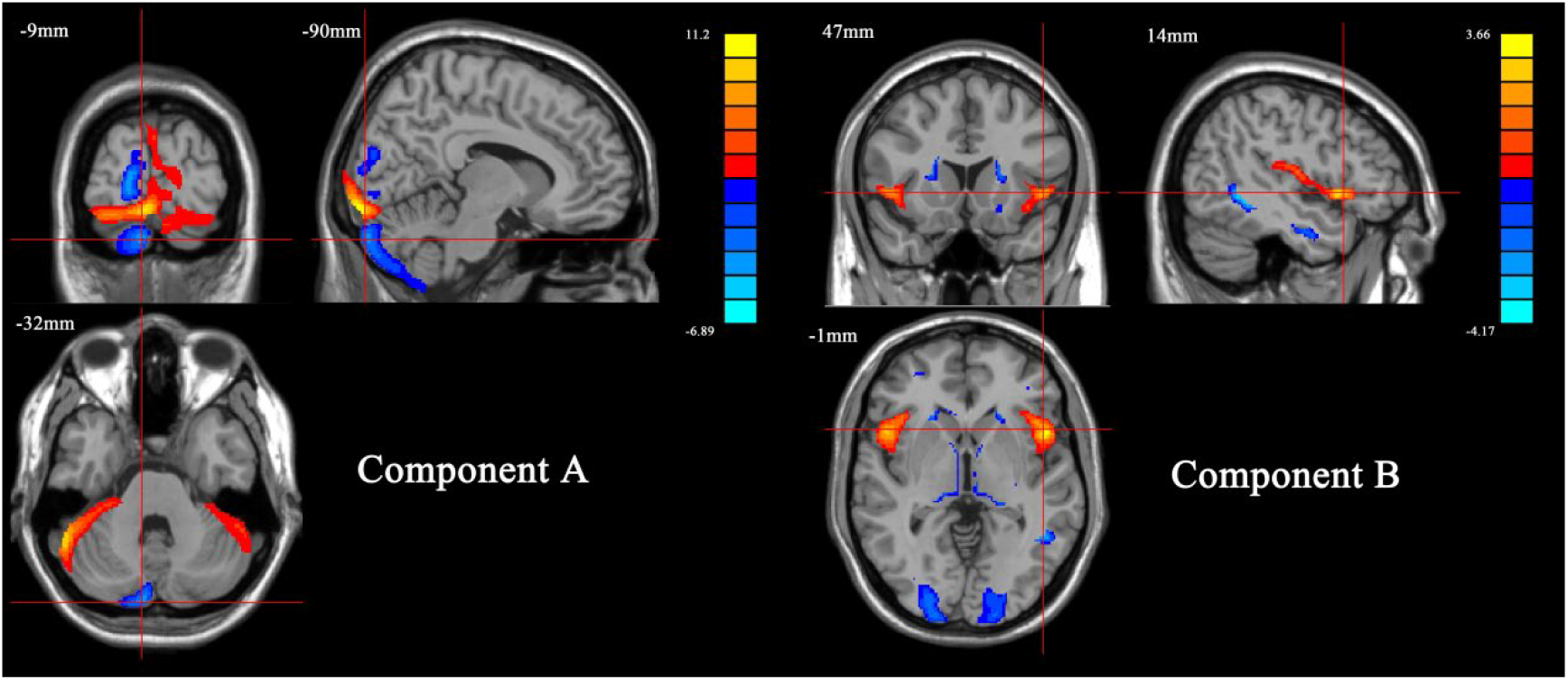
The two components showed significant case/control difference. Component A mainly included bilateral Crus I, left lingual gyrus, left Crus II, and left fusiform (Z score > 2, cluster volume > 1.5 cc^3^). Component B mainly included bilateral insula, caudate, thalamus and middle occipital gyrus (Z score > 2, cluster volume > 1.5 cc^3^).

**Table 1.** Demographics of the sample

|  | ADHD (210) | Unaffected<br>siblings (108) | Subthreshold<br>Subjects (49) | Controls (141) |
| --- | --- | --- | --- | --- |
| Age (years) | 14.61 ± 2.41 | 14.59 ± 2.16 | 15.13 ± 1.97 | 14.58 ± 2.14 |
| Gender (F/M) | 81/129 | 68/40 | 16/33 | 62/79 |
| Estimated IQ | 98.63 ± 16.49<br>(0.5%) | 99.33 ± 14.15<br>(0%) | 98.79 ± 12.42<br>(0%) | 105.13 ± 13.91<br>(1.4%) |
| Inattention | 7.33 ± 1.71<br>(0%) | 0.44 ± 1.11<br>(0%) | 4.34 ± 0.95<br>(0%) | 0.33 ± 1.11<br>(0%) |
| Hyperactivity<br>/Impulsiveness | 5.99 ± 1.71<br>(0%) | 0.34 ± 0.79<br>(0%) | 2.73 ± 1.96<br>(0%) | 0.17 ± 0.63<br>(0%) |
| Maximum digit<br>span forward | 7.94 ± 1.79<br>(0.5%) | 8.62 ± 1.70<br>(0%) | 8.80 ± 1.79<br>(0%) | 8.70 ± 1.68<br>(1.4%) |
| Maximum digit<br>span backward | 5.10 ± 1.74<br>(0.5%) | 6.06 ± 1.67<br>(0%) | 6.06 ± 2.10<br>(0%) | 6.05 ± 1.94<br>(1.4%) |
| Stop task<br>errors | 9.45 ± 12.1<br>(40%) | 5.64 ± 8.02<br>(36%) | 5.47 ± 4.33<br>(39%) | 4.47 ± 6.76<br>(31%) |
| Stop task<br>SSRT | 277.79 ±<br>76.01<br>(40.5%) | 256.05 ± 8.02<br>(36%) | 282.42 ± 58.29<br>(39%) | 263.14 ± 6.76<br>(31%) |
| Stop task<br>ICV | 0.22 ± 0.05<br>(40%) | 0.19 ± 0.04<br>(36%) | 0.20 ± 0.05<br>(39%) | 0.18 ± 0.04<br>(31%) |
| SWM average<br>accuracy | 0.67 ± 0.15<br>(42%) | 0.68 ± 0.15<br>(44%) | 0.74 ± 0.11<br>(31%) | 0.74 ± 0.041<br>(23%) |
| Scan site 1 | 123 | 52 | 22 | 46 |
| Scan site 2 | 87 | 56 | 27 | 95 |

Note: Values displayed in the table were showed as mean ± standard deviation. Site 1 scanner was AVANTO, and site 2 was SONATA. The percentages in parentheses show the missingness of data in each diagnostic group. Stop task error includes N commission and omission errors together. Stop task SSRT: stop task stop signal reaction time. Stop task ICV: interindividual coefficient of variation which is the reaction time variance/mean reaction time. SWM average accuracy: the average accuracy of visuo-spatial working memory task.

Two components showed significant case/control difference passing FDR correction, consistently showing lower loading coefficients in cases than controls. Component A (corrected *p* = 0.00029; Cohen’s *d* = −0.32) was mainly located in the bilateral Crus I while controlling for age (*p* = 7.85e-13; β = −0.15) and medication (*p* = 0.00070; β = 0.41) in the model. Component B (corrected *p* = 0.027, Cohen’s *d* = −0.31) was found largely in the bilateral insula while controlling for age (*p* = 0.046; β = 0.40), age^2 (*p* = 0.0051; β = −0.020), and medication (*p* = 0.041; β = 0.24) in the model. See Figure 1 and Table 2 for detailed representations of these two components. The moving average of loading coefficients of component A and B are showed in Figure 2.

**Figure 2.**
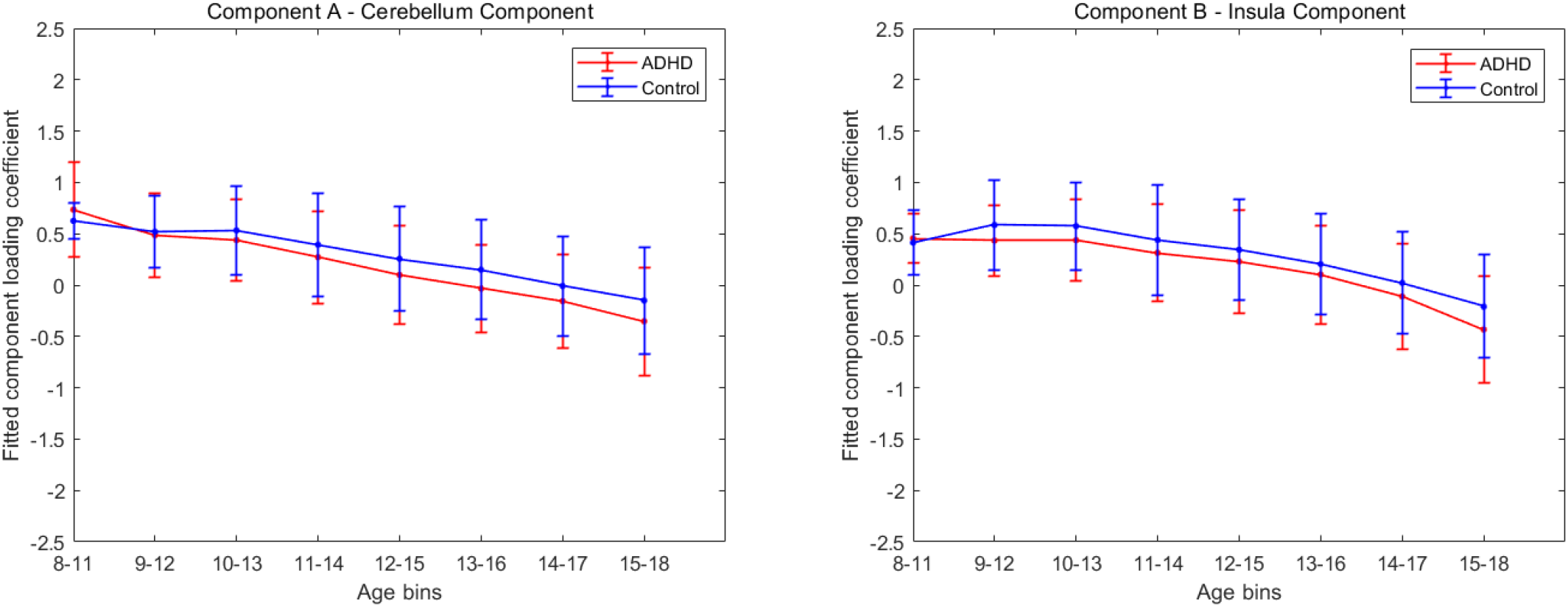
Moving averages of component A loading coefficients, corrected for age, medication and family structures are showed on the left. Those of component B loadings coefficients, corrected for age, age^2, medication, and family structures are showed on the right. Age bins with less than 5 subjects in either group are removed from the figures.

**Table 2.** Brain regions, peak coordinates and volumes of components A and B.

|  | Brain regions | Volumes<br>(cc<sup>3</sup>) | Peak coordinates<br>(Z and coordinates) |
| --- | --- | --- | --- |
| Component A |  |  |  |
| Cluster 1 | Crus I (extended to cerebellum posterior lobe and L lingual gyrus) | 2.1 | 11.2 (-9, -95, -14) |
| Cluster 2 | L Crus II | 0.9 | -4.7 (-9, -90, -32) |
| Cluster 3 | L fusiform gyrus | 0.3 | -6.8 (-29, -75, -3) |
| Component B |  |  |  |
| Cluster 1/2 | L/R insula (extended to inferior frontal gyrus) | 0.7/0.7 | 3.3(-50, 12, -3)/3.7 (48, 12, 3) |
| Cluster 3/4 | L/R middle occipital gyrus | 0.8/1.0 | -4.2(-33,-74,15)/-4.2(30, -66, 25) |
| Cluster 5 | L/R caudate | 0.3/0.4 | -2.9(-21,23,7)/-2.7(19,22,7) |
|  | L/R thalamus | 0.3/0.3 | -3.1(-16,25,3)/-2.8(9,-19,3) |
|  | R inferior temporal gyrus | 0.5 | -4.2(51, -50, -6) |

Note: For both components, Z score was set $> 2$ , and cluster volumes were set $> 1.5 \text{ cc}^3$ to retain their most significant results. The directions of peak Z scores indicate whether the brain region contributed positively or negatively to the component as red for positive and blue for negative in brain maps.

Component A showed significant differences only in the comparison between cases and controls; sub-clinical cases on average were intermediate between cases and unaffected siblings, who were not significantly different from controls. In component B, the unaffected siblings also showed significantly reduced loadings relative to controls (corrected *p* = 0.04), similar to cases, while subthreshold cases did not (see Figure 3).

**Figure 3.**
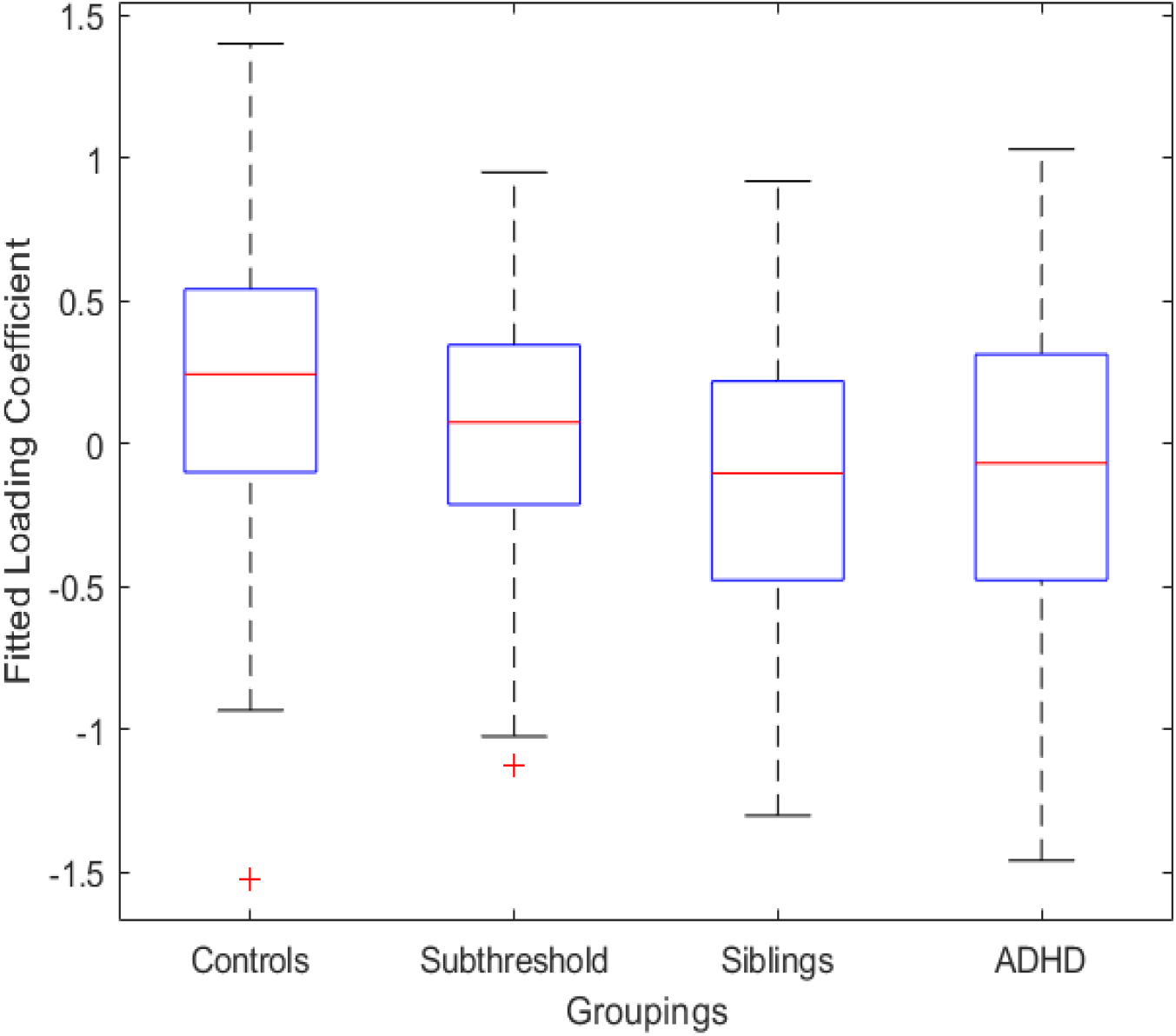
In component B, persons with ADHD showed significant reduced loading compared to controls (corrected *p* = 0.027, Cohen’s d = −0.31), and the unaffected siblings also showed significantly reduced loadings relative to controls (corrected *p* = 0.04), similar to cases while subthreshold cases did not. The whiskers extend to the most extreme data points not considered outliers, and the outliers are plotted individually using the ‘+’ symbol.

No association was found between both components’ loadings and performance on any of the cognitive tests. However, after correcting for medication and IQ, component A showed a negative correlation with inattention symptoms across the entire group (corrected *p* = 0.011, β = −0.43, VE = 1.4%; see Figure 4).

**Figure 4.**
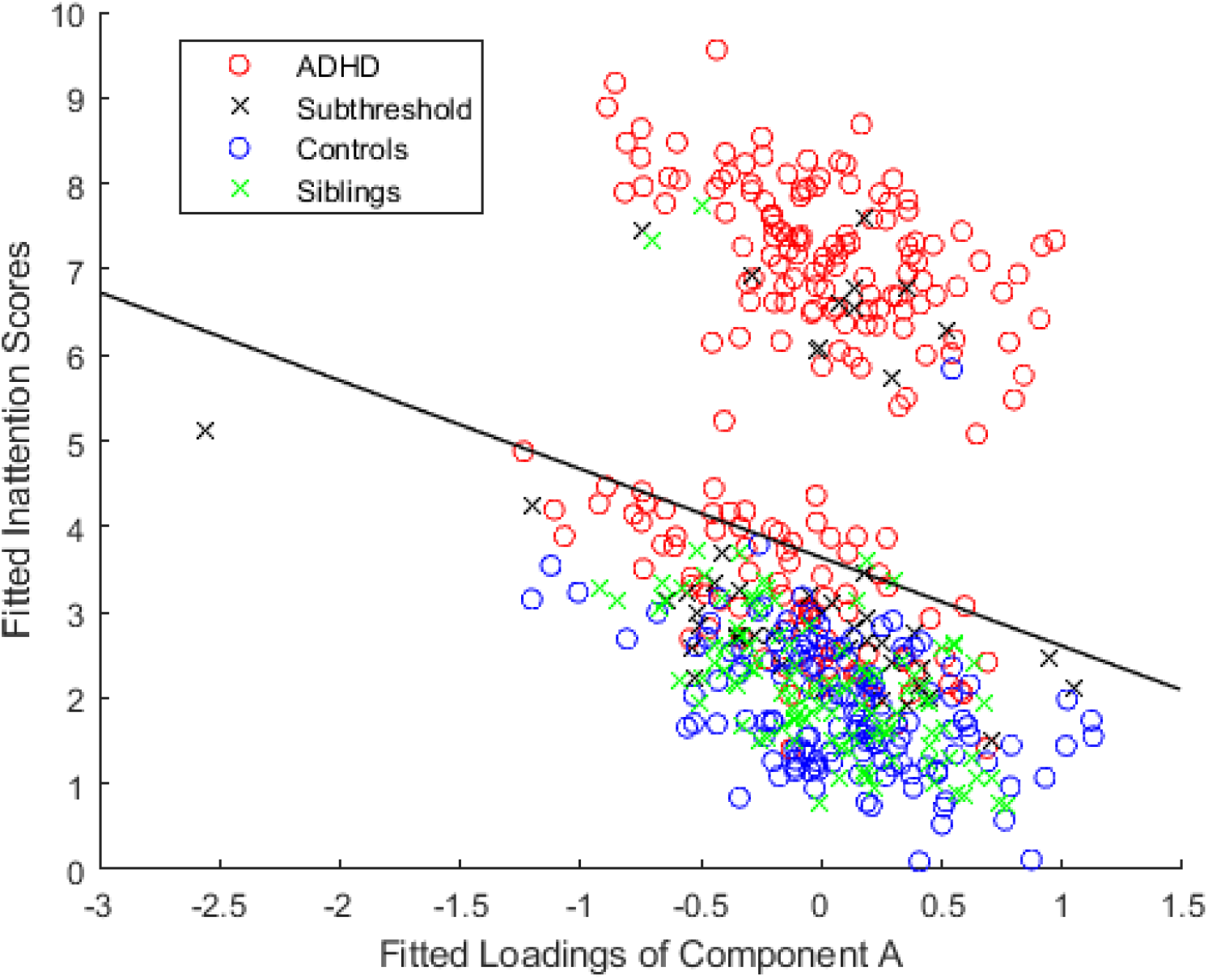
Component A showed a negative correlation with inattention symptoms across the entire group (corrected *p* = 0.011, beta = −0.43, VE = 1.4%) after correcting for IQ and medication. In the figure, fitted values of loading coefficient and inattention scores from the linear mixed model were plotted.

## 4. Discussion

In this study, we used an SBM analysis to identify gray matter network differences in individuals with ADHD, unaffected siblings, sub-clinical ADHD participants, and unrelated healthy controls. Using a hypothesis free approach (no pre-selection of regions of interest), we identified two components, in the cerebellum and insula, that showed significant gray matter reductions in participants with ADHD compared to controls. We also found this reduction continued through the whole adolescence, similar to previous findings using the same age range in a larger cohort that included the current study’s participants (Hoogman et al., 2017). In addition, unaffected siblings exhibited gray matter loadings similar to controls in the cerebellar component (component A), but showed a negative relationship to inattention levels independent of clinical status. The insula component (component B), in contrast, showed a reduction in unaffected siblings similar to that in cases, but no significant relationship was observed with symptom severity or cognitive performance. In addition, there were no components significantly related to any of the cognitive measures. Overall, this approach suggests that brain areas reflecting genetic liability within ADHD may be partly separate from areas modulating symptom severity, as has been suggested previously (van Ewijk et al., 2014; Wu et al., 2017).

In component A, the bilateral Crus I (extending to cerebellum posterior lobe and left lingual gyrus) contributed most strongly, and the left Crus II and left fusiform also contributed, though less strongly, to this component. The composition of this component is consistent with the diverse functions of the cerebellum. Of particular relevance to ADHD, functions of the cerebellum include motor control, working memory, and attention (Duan et al., 2018; Ivanov et al., 2014; Moore et al., 2017). Multiple studies now suggest that the cerebellum, and alterations in its structure and function, play an important role in ADHD throughout childhood, adolescence, and adulthood (Duan et al., 2018; Noordermeer et al., 2016; Valera et al., 2007). Our analyses support the current literature, with a focus on Crus I and Crus II, which are considered part of an executive control network (Stoodley, 2012). Executive function deficits are known to be a key problem in ADHD (Mahone and Denckla, 2017; Mueller et al., 2017) and have been related to inattention symptoms (Chhabildas et al., 2001; Neely et al., 2016; Nigg et al., 2005; Willcutt et al., 2005).

The role of the cerebellum is backed by previous analyses in an overlapping data set of the NeuroIMAGE project. A linked ICA analysis of both children and adult brain images, taking into account fractional anisotropy, mean diffusivity, diffusion mode, VBM, cortical thickness estimate and area expansion estimate, reported reduced Crus I, Crus II, cerebellar tonsil and culmen volume in its VBM component in ADHD compared to non-ADHD participants (Francx et al., 2016). Multivariate analysis in adults with ADHD also showed a brain component involving cerebellar tonsil and culmen, and their volume negatively correlated with inattention scores (the less volume worse, the inattention symptoms), while a voxel-wise analysis showed cerebellum gray matter volume positively correlated with working memory in the same study (Duan et al., 2018). Our findings in component A overlap with these previous findings of the cerebellum showing reductions in ADHD compared to non-ADHD, though the cognitive relationship in our analysis was inattention symptoms rather than working memory performance.

Component B also showed gray matter reduction compared to the controls, and it mainly included the insula, extending to inferior frontal gyrus, and included bilateral caudate, bilateral thalamus, right inferior temporal gyrus and bilateral middle occipital gyrus. Functional neuroimaging studies have emphasized the key role of the insula during salience processing and inhibitory control, with ADHD patients showing abnormal activation/deactivation (Cubillo et al., 2012; Rubia et al., 2011) or hyperactivation in the insula during distracting stimuli (Vetter et al., 2018). In contrast, the reduction of gray matter volume of the insula has been previously reported in ADHD, (Lopez-Larson et al., 2012) as well as in common ADHD comorbidities of inhibitory control, including obsessive-compulsive disorder (Norman et al., 2016) and oppositional defiant disorder (Noordermeer et al., 2017). In this context of similar disorders, the insula has often been implicated in larger inhibitory control networks including ventrolateral prefrontal cortex, supplementary motor area, dorsal anterior cingulate cortex, and the striatum, thalamus, and parietal regions (Aron, 2011; Hugdahl et al., 2015). However, in this structural analysis, we did not find a relationship with inattention or other cognitive or symptom measures for this network.

Involvement of subcortical structures in component B is consistent in with findings from large-scale studies of ADHD. The largest subcortical study to date, from the ENIGMA ADHD group, showed that most subcortical regions, including nucleus accumbens, amygdala, caudate nucleus, hippocampus, and putamen were smaller in ADHD cases than controls (Hoogman et al., 2017). However, none of the previous univariate meta-analysis of VBM results have reported the significance of the bilateral insula, despite its important role in inhibitory control. Our SBM findings for component B provide a more insightful profile of how different brain regions may stably work together to form a recognizable network than if we had investigated their volume alterations individually. Finally, component B might be capturing part of the genetic loading involved in brain development shared by ADHD patients and their unaffected siblings, independent of symptom severity and cognitive performance. Although these two components were found through ICA; connectivity between the cerebellum and the insula has been documented in both resting state and functional studies, showing support for potential biological connections between our findings (Balsters et al., 2014; Buckner et al., 2011).

Strengths of this study include multivariate analyses on participants from both ADHD and control families with family structures taken into account in the linear mixed model. In addition, the relationship of brain components among categories (ADHD, unaffected siblings, sub-clinical participants and controls) and symptom dimensions were also examined. The previous multivariate analysis in the NeuroIMAGE project using linked ICA was concordant with our work by showing reduced cortical thickness in the insula, occipital lobes and anterior cingulate compared to the non-ADHD participants (Francx et al., 2016). The VBM analysis from the NeuroIMAGE project had shown, in addition to the previously reported prefrontal, inferior frontal and occipital regions (Durston et al., 2004), that ADHD patients had cortical deficits in the precentral gyrus, medial and orbitofrontal cortex, and cingulate gyrus; while their unaffected siblings showed the same differences excluding the precentral gyrus (Bralten et al., 2016). However, multivariate analyses identify complementary brain deficits to these cortical alterations. The cerebellum, insula and other regions also played a role, and these alterations have been previously shown to group together (Duan et al., 2018; Francx et al., 2016).

These results help distinguish symptom dimensions from genetic liability. The cerebellum component extracted by SBM was more closely connected to inattention symptoms across the four groups, and it was not strongly affected by the common genetic factors shared in family structures. The insula component captured the alterations shared by ADHD and their siblings, but no component was related to the symptom dimensions. This contrast might suggest different mechanisms were contributing to the complicated ADHD clinical profile, which was also discovered in previous studies (van Ewijk et al., 2014; Wu et al., 2017).

A limitation was that age, gender, and scanning site were not well-balanced in each group as a result of original data collection limitations. The sample also covered a relatively broad age span from 7 to 18 years, which might bring in potential brain development bias. To counteract such limitations, we did a voxel-wise correction for gender and site, and examined whether age or age^2 showed a significant influence on the brain images for each component (in their respective models). In addition, the stop task did not have data available for all participants, which might have lowered the power for detecting associations between the gray matter and inhibition.

## 5. Conclusion

In conclusion, our data suggest that the cerebellum and insula components might shed light on different but related mechanisms of ADHD, indicating that the clinical phenotype, cognitive performance, and mechanisms shared in family structure can be supported by different brain networks. Gray matter abnormalities were found to either underlie inattention symptoms or be affected by shared relationships between ADHD patients and their unaffected siblings. This suggests that brain areas reflecting genetic liability within ADHD may be partly separate from areas modulating symptom severity.

## Author contributions

J. Turner, J. Liu and W. Jiang designed the study. A. Arias-Vasquez acquired the data and consulted on the interpretation. W. Jiang and K. Duan analyzed the data. W. Jiang and K. Rootes-Murdy wrote the article, which all authors reviewed. All authors approved the final version to be published and can certify that no other individuals not listed as authors have made substantial contributions to the paper.

## Acknowledgments

This study was supported by the National Institutes of Health and The National Institute of Mental Health through the grant 1R01MH106655. This NeuroIMAGE study was supported by NIH Grant R01MH62873, NWO Large Investment Grant 1750102007010 and grants from Radboud University Medical Center, University Medical Center Groningen and Accare, and VU University Amsterdam. This work was also supported by grants from NWO Brain & Cognition (433-09-242 and 056-13-015) and from ZonMW (60-60600-97-193). Further support was received from the European Union’s FP7 program under grant agreement no. 278948 (TACTICS), no. 602450 (IMAGEMEND), no. 602805 (Aggressotype), and from the European Union’s Horizon 2020 research and innovation program under grant agreement no. 667302 (CoCA) and no. 728018 (Eat2beNICE). Barbara Franke receives funding from a personal Vici grant (to Barbara Franke) of the Netherlands Organization for Scientific Research (NWO, grant numbers 433-09-229 and 016-130-669) and a pilot grant of the Dutch National Research Agenda for the NeuroLabNL project.

