## Supplemental Figure 1 for "Structural Brain Alterations and Their Association with Cognitive Function and Symptoms in Attention-Deficit/Hyperactivity Disorder Families"

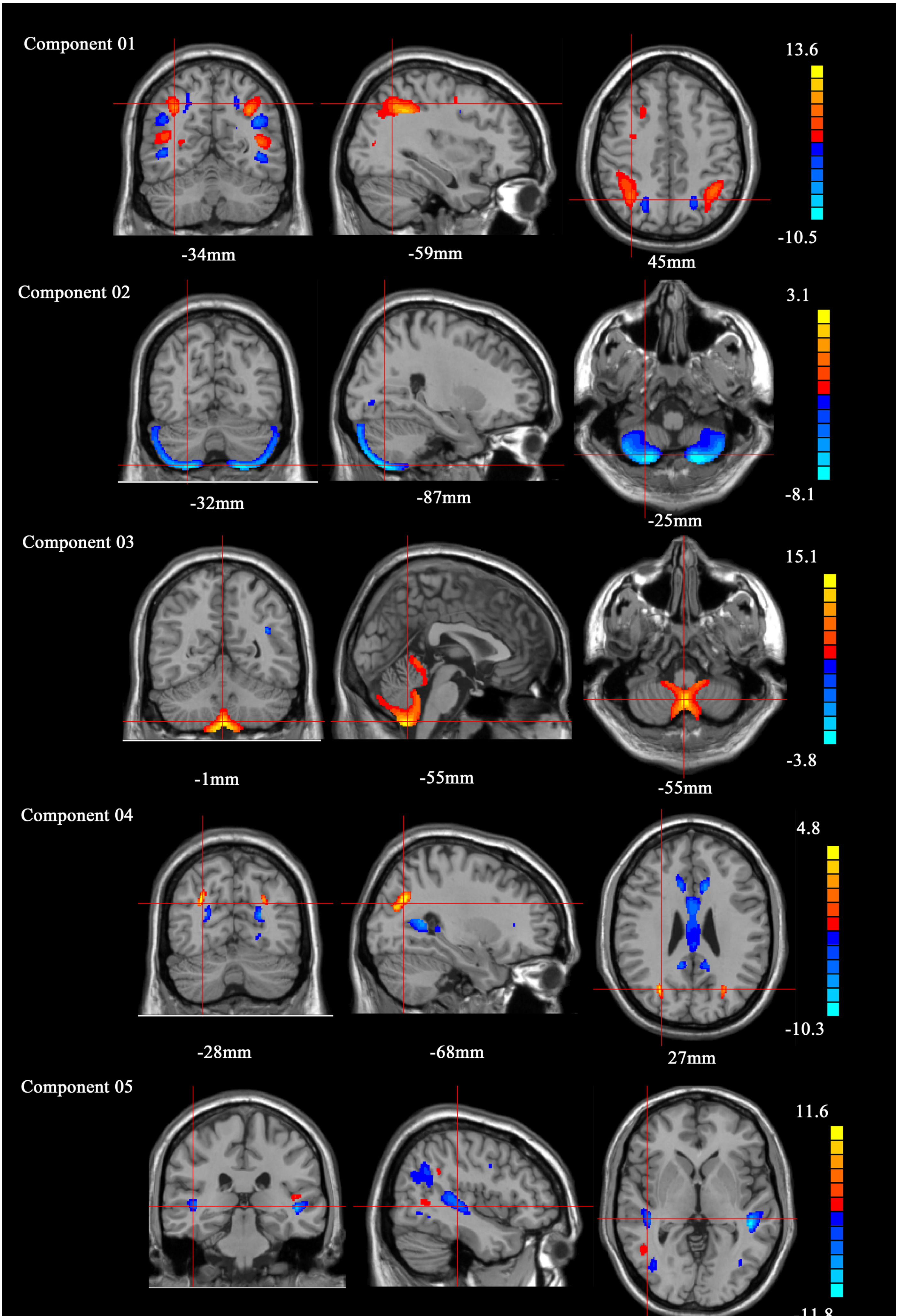
Component 01
13.6
-10.5
-34mm
-59mm
45mm
Component 02
3.1
-8.1
-32mm
-87mm
-25mm
Component 03
15.1
-3.8
-1mm
-55mm
-55mm
Component 04
4.8
-10.3
-28mm
-68mm
27mm
Component 05
11.6

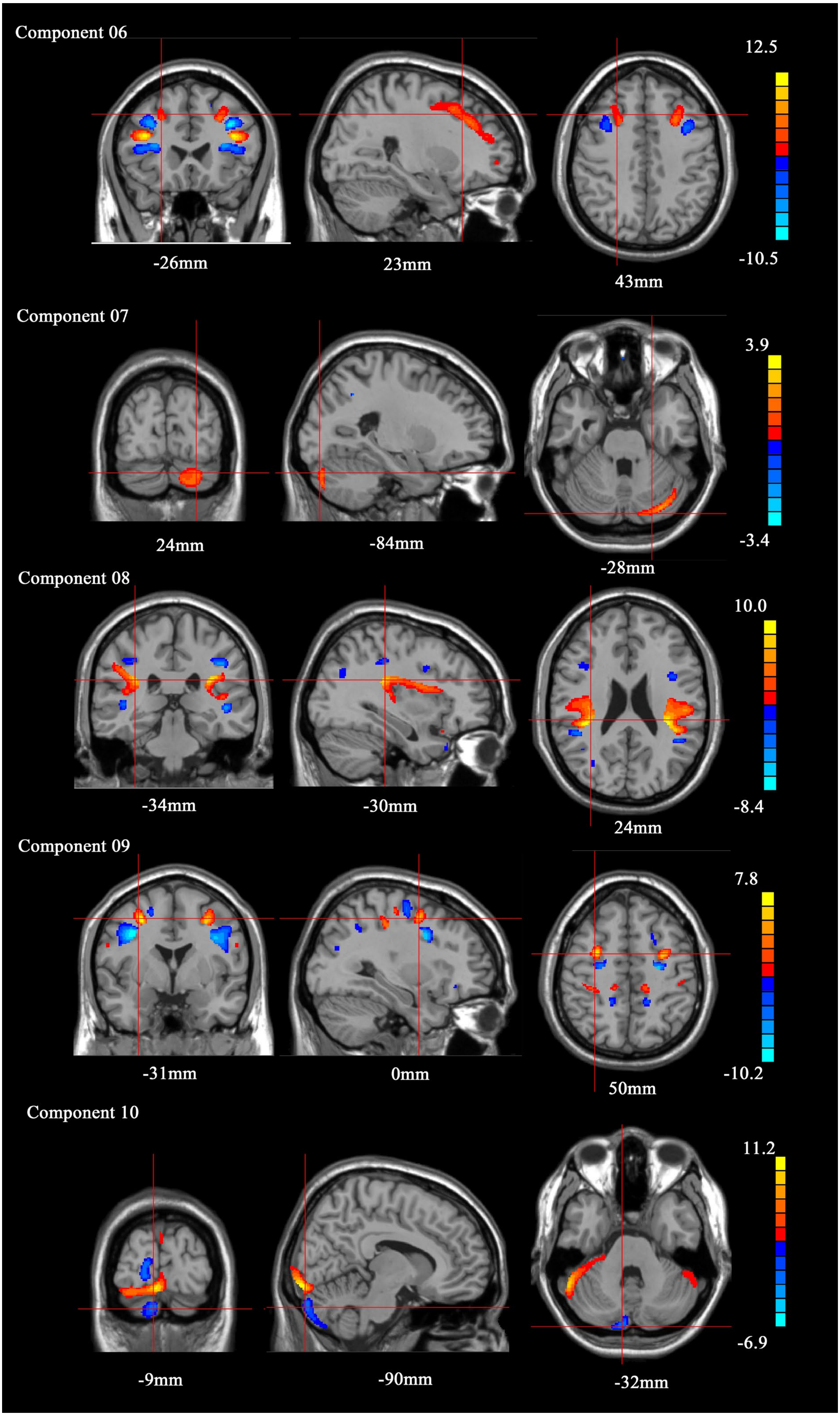
Component 06
-26mm
23mm
43mm
12.5
-10.5
Component 07
24mm
-84mm
-28mm
3.9
-3.4
Component 08
-34mm
-30mm
24mm
10.0
-8.4
Component 09
-31mm
0mm
50mm
7.8
-10.2
Component 10
-9mm
-90mm
-32mm
11.2
-6.9

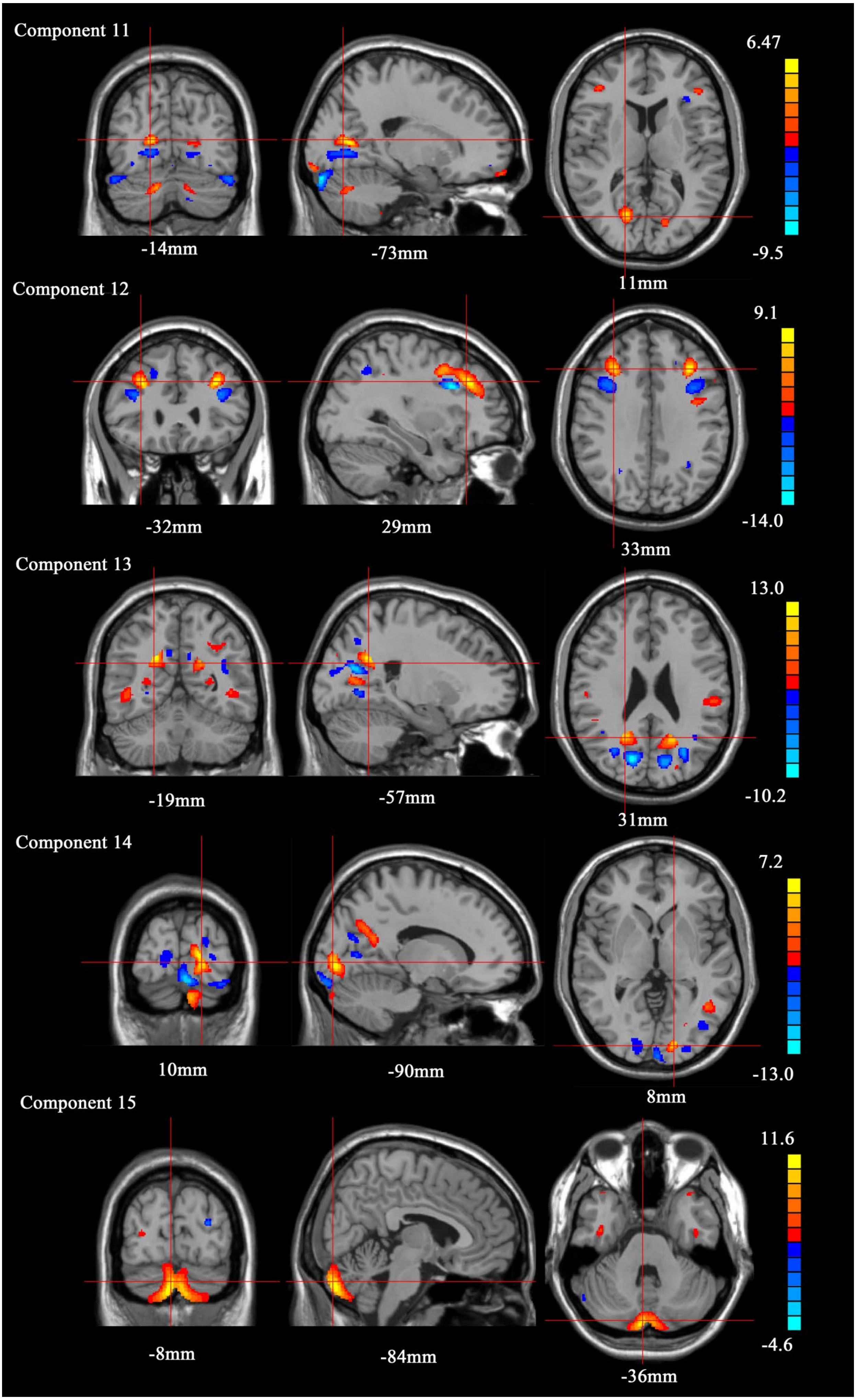

Component 11
6.47
-9.5
-14mm
-73mm
11mm
Component 12
9.1
-14.0
-32mm
29mm
33mm
Component 13
13.0
-10.2
-19mm
-57mm
31mm
Component 14
7.2
-13.0
10mm
-90mm
8mm
Component 15
11.6
-4.6
-8mm
-84mm
-36mm

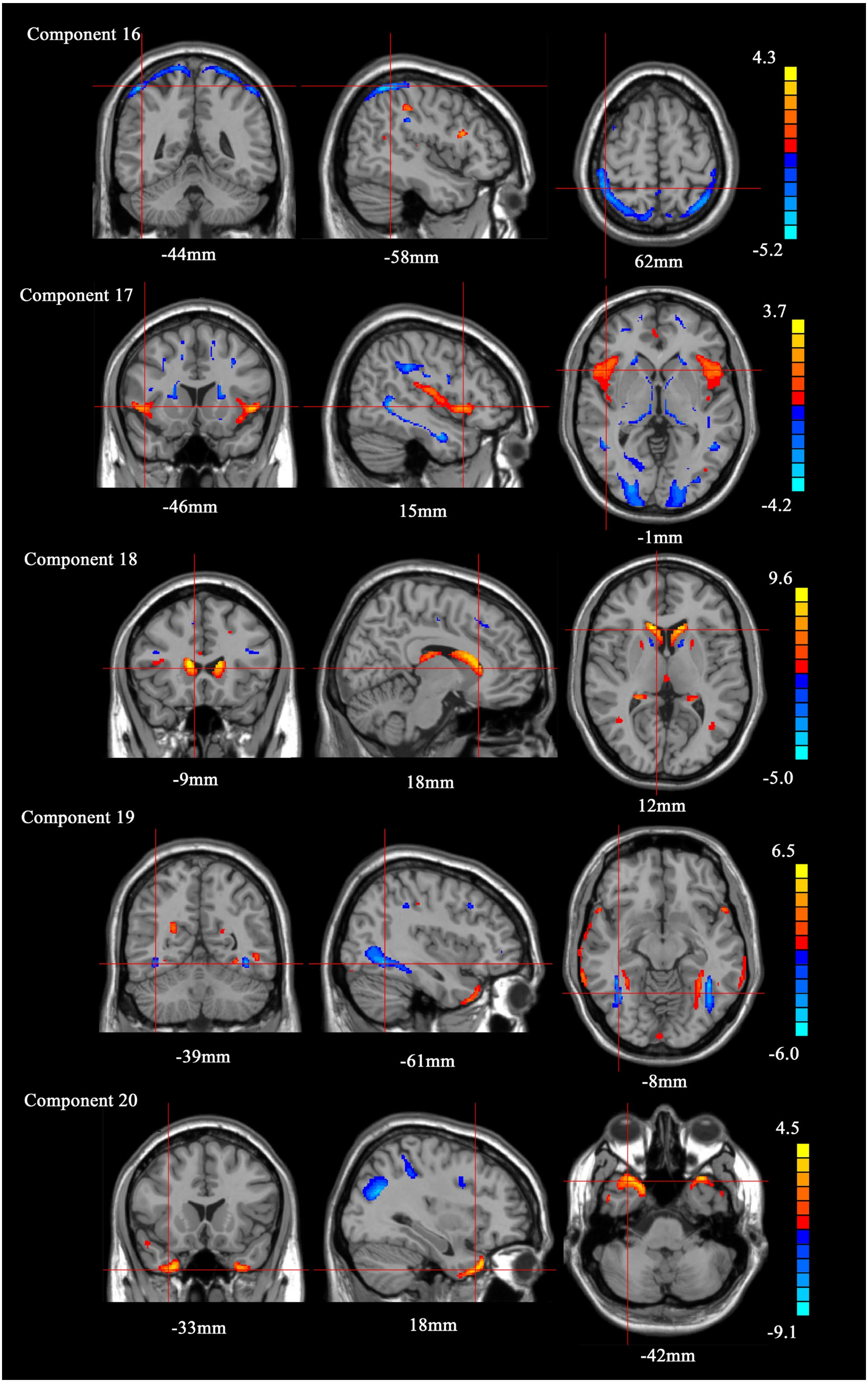
Component 16
-44mm
-58mm
62mm
4.3
-5.2
Component 17
-46mm
15mm
-1mm
3.7
-4.2
Component 18
-9mm
18mm
12mm
9.6
-5.0
Component 19
-39mm
-61mm
-8mm
6.5
-6.0
Component 20
-33mm
18mm
-42mm
4.5
-9.1

Supplemental figure 1. The T1 images were decomposed into 20 stable components (all set to Z score > 2), and in the main text component A was component 10 while component B was component 17.
